# Development of a Cx46 targeting strategy for cancer stem cells

**DOI:** 10.1101/289397

**Authors:** Erin E. Mulkearns-Hubert, Luke A. Torre-Healy, Daniel J. Silver, Jennifer T. Eurich, Emily Serbinowski, Masahiro Hitomi, John Zhou, Bartlomiej Przychodzen, Renliang Zhang, Samuel A. Sprowls, James S. Hale, Tyler Alban, Artem Berezovsky, Brent A. Bell, Paul R. Lockman, Babal K. Jha, Justin D. Lathia

## Abstract

Gap junction-mediated cell-cell communication enables tumor cells to synchronize the execution of complex processes. Despite the connexin family of gap junction proteins being considered tumor suppressors, we previously found that glioblastoma cancer stem cells (CSCs) express higher levels of Cx46 compared to non-stem tumor cells, and this was necessary and sufficient for CSC maintenance. To develop a Cx46 targeting strategy, we utilized point mutants to disrupt specific functions of Cx46 and found that gap junction coupling was the critical function of Cx46 for CSCs. Based on this finding, we screened a clinically relevant library of small molecules and identified clofazimine as an inhibitor of Cx46-specific cell-cell communication. Clofazimine attenuated proliferation, self-renewal, and tumor growth and synergized with temozolomide to induce apoptosis. These data suggest that combining clofazimine with standard-of-care therapies may target glioblastoma CSCs. Furthermore, these results demonstrate the importance of targeting cell-cell communication as an anti-cancer therapy.

## Introduction

Glioblastoma (GBM; grade IV astrocytoma), the most commonly occurring primary malignant brain tumor, remains uniformly fatal despite aggressive therapy including surgery, radiation, and chemotherapy. Research advances have increased the understanding of the disease and identified new therapies, but patient prognosis remains poor, with a median survival of only 14-16 months, and 5-year-survival rates remain less than 3% (McGirt et al., 2009; Stupp et al., 2009; Stupp et al., 2015). One factor underlying the difficulty in treating GBM is the cellular diversity present within these tumors. Heterogeneous populations of cancer stem cells (CSCs) exhibit essential characteristics of sustained self-renewal, persistent proliferation, and the ability to initiate tumors if transplanted into mice (Lathia et al., 2015) and also display resistance to the GBM standard-of-care therapies radiation and temozolomide (Bao et al., 2006; Chen et al., 2012; Liu et al., 2006). Current efforts to treat GBM are focused on the ability to target CSCs, as this may lead to the development of more effective therapies for GBM with increased clinical success.

Cell-cell communication is mediated through the connexin family of proteins and the gap junction (GJ) channels these proteins comprise. Six connexin proteins assemble into a channel through the plasma membrane that can exchange small molecules between the cell and the extracellular space as hemichannels. When these channels dock with a compatible hexamer on a neighboring cell, a GJ is formed. GJ intercellular communication (GJIC) exchanges ions, miRNAs, small metabolites such as glucose, antioxidants, and peptides between cells, allowing them to coordinate their phenotypes and respond to environmental conditions (Goodenough and Paul, 2009). Connexin proteins serve three main cellular functions: exchange of small molecules between cells as GJs, exchange of small molecules between a cell and the extracellular space as hemichannels, and intracellular protein-protein interactions (Goodenough and Paul, 2003, 2009; Leithe et al., 2018; Stout et al., 2004).

Previous work based mainly on connexin 43 (Cx43) suggested that connexins act as tumor suppressors (Aasen et al., 2016). However, we have identified pro-tumorigenic connexins in prostate cancer (Zhang et al., 2015), breast cancer (Thiagarajan et al., 2018), leukemia (Sinyuk et al., 2015), and GBM (Hitomi et al., 2015). GBM CSCs express higher levels of Cx46 compared to non-stem tumor cells (non-CSCs), and Cx46 is required for CSC proliferation, survival, self-renewal, and tumor formation (Hitomi et al., 2015). Pan-gap junction inhibitors slowed tumor growth in mice with intracranial tumors, but these compounds inhibit connexins as an off-target effect. For this reason, these compounds would likely cause side effects in patients based on their broad effects targeting multiple connexins that play essential roles in many normal organs. Here, we used mutational analysis and identified the dominant function of Cx46 in GBM CSCs to be cell-cell communication through gap junctions (GJIC) rather than hemichannel activity. We thus hypothesized that targeting of CSCs through specific inhibition of Cx46 would slow tumor growth and lead to the development of new therapies for patients with GBM. A screen of FDA-approved small molecules identified the anti-leprosy drug clofazimine as a preferential inhibitor of Cx46-mediated cell-cell communication and GBM CSC maintenance, suggesting that repurposing of this and similar compounds may benefit patients with GBM.

## Results

### Cx46-mediated cell-cell communication is essential to maintain glioblastoma cancer stem cells

Our previous studies indentified Cx46 as a potential anti-CSC target. To develop a strategy to inhibit Cx46, we first sought to determine the function of Cx46 required to maintain GBM CSC properties. To achieve this, we identified a panel of Cx46 mutations that would allow us to deduce the individual importance of GJIC and hemichannel activity. Two Cx46 point mutations have been reported in human patients with cataracts (Hansen et al., 2006; Santhiya et al., 2010). These mutations, L11S and T19M, are both located in the N-terminal tail of the Cx46 protein (**Fig. 1A**) and have been functionally investigated in the context of the rat protein in *Xenopus* oocytes (Tong et al., 2015; Tong et al., 2013). When co-expressed with wild-type Cx46, the presence of the L11S mutation dramatically reduced both GJIC and hemichannel activity (Tong et al., 2013). In contrast, co-expression of the Cx46 T19M mutant with wild-type Cx46 increased hemichannel activity but did not affect GJIC (Tong et al., 2015). We also utilized a “cysless” mutant previously engineered in Cx43 that disrupts the three disulfide bonds necessary to maintain the structure of connexins required for gap junction docking. This mutant was reported to completely block GJIC without affecting hemichannel activity of Cx43 in both *Xenopus* oocytes and ovarian granulosa cells (Bao et al., 2004; Tong et al., 2007).

**Figure 1.**
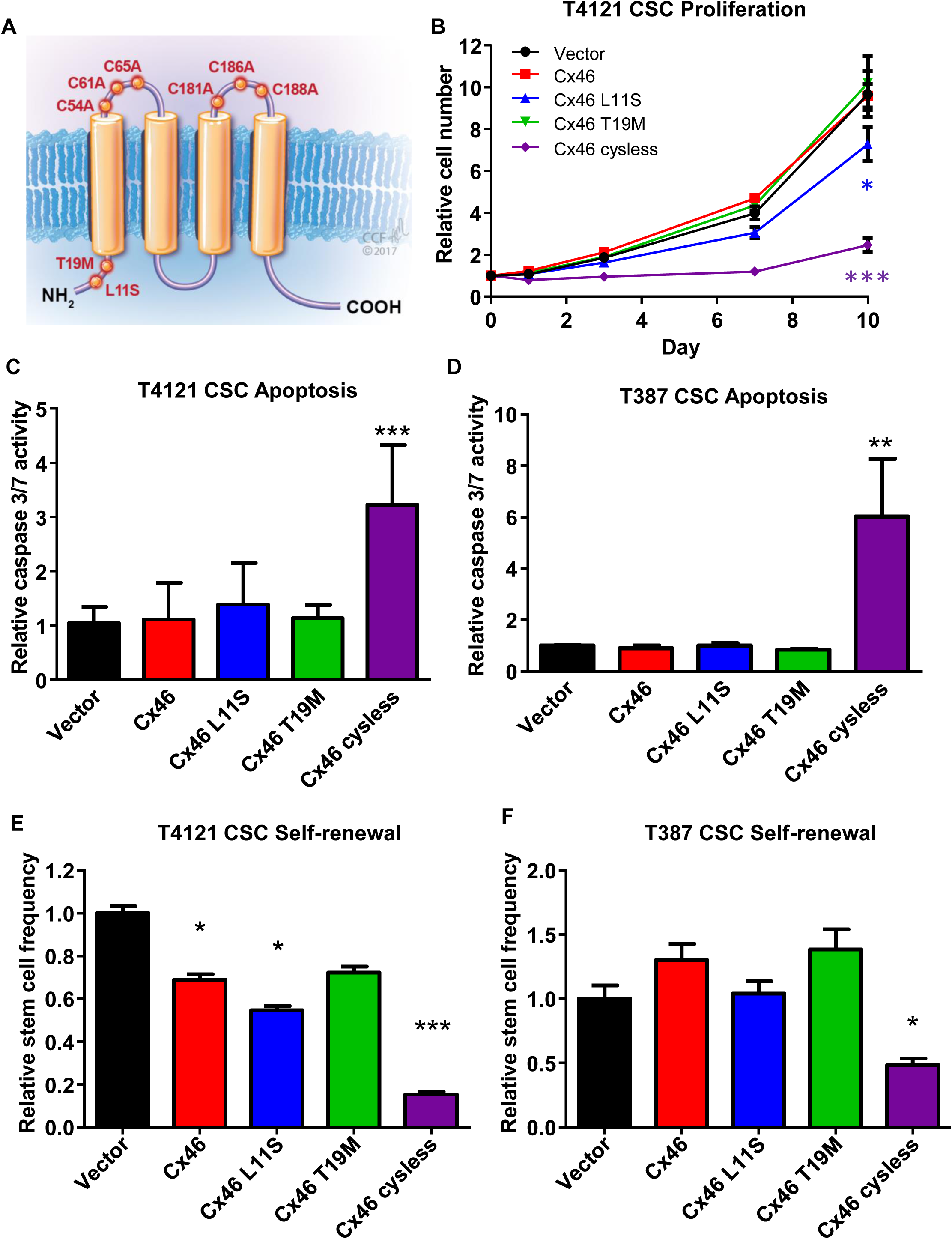
Mutational analysis indicates that cell-cell communication is essential to maintain glioblastoma cancer stem cells. (**A**) Schematic showing the location of Cx46 point mutants in the protein. (**B**) CSCs from the patient-derived xenograft specimen T4121 were transfected with wildtype or mutant Cx46, and the number of cells was measured on days 0, 1, 3, 7, and 10 after plating using CellTiter-Glo. The values shown are relative to day 0. n = 4 experiments performed in triplicate. * p<0.05, ** p<0.01, *** p<0.001 by two-way ANOVA compared to vector to test for significant differences between the curves. (**C-D**) Transfected CSCs from the patient-derived xenograft specimens T4121 (**C**) and T387 (**D**) were assessed for active caspase 3/7 on day 1 using Caspase-Glo. The values shown are normalized to the CellTiter-Glo signal at day 1, and the values shown are given relative to vector. n = 4 experiments for T4121 and n = 3 for T387, all performed in triplicate. * p<0.05, ** p<0.01, *** p<0.001 by Student’s unpaired t-test with Welch’s correction compared to vector. (**E-F**) Transfected CSCs from the patient-derived xenograft specimens T4121 (**E**) and T387 (**F**) were plated in a limiting-dilution format (between 1-20 cells/well of a 96-well plate), and the number of spheres per well was counted between days 10 and 14. The stem cell frequency was calculated using the online algorithm described in the Methods section. The values shown are relative to the stem cell frequency of the vector-transfected cells. n = 3 experiments for T4121 and n = 2 for T387, with 24 technical replicates per cell number per experiment. * p<0.05, ** p<0.01, *** p<0.001 by χ^2^ test compared to the vector control. Data are represented as mean ± SEM for B-D and mean ± range for E-F. See also Supplemental Figure S1.

We introduced these mutations into human Cx46 cDNA and transfected GBM CSCs isolated from two different patient-derived xenografts (T4121 and T387) with the three constructs. Using qPCR, we were able to detect the expression of each Cx46 mutant in CSCs at the mRNA level (**Supplemental Fig. S1A-B**). Expression of the Cx46 T19M mutant or overexpression of wild-type Cx46 had little effect on CSC proliferation, apoptosis, or self-renewal, a hallmark of the CSC state, which was assessed by limiting dilution sphere-formation analysis (**Fig. 1B-F, Supplemental Fig. S1C**), while we observed slight decreases in proliferation and self-renewal with expression of Cx46 L11S. However, expression of the Cx46 cysless mutant induced a decrease in CSC proliferation, an increase in apoptosis, and a decrease in self-renewal in both patient-derived specimens. This observation demonstrates that the cysless mutant, which has been shown to have the greatest effect on cell-cell communication, also has the greatest effect on CSC maintenance (**Supplemental Fig. 1D**) and led us to conclude that GJIC mediated by Cx46 is essential to maintain GBM CSC proliferation, survival, and self-renewal.

### A screen of FDA-approved small molecules identifies clofazimine as an inhibitor of Cx46-mediated cell-cell communication

Based on our observation that GBM CSCs require Cx46-mediated GJIC for survival, we designed an assay system to screen for inhibitors of this process. We assessed GJIC using a quantitative calcein transfer assay (**Fig. 2A**) (Hitomi et al., 2015), a modification of the parachute dye-uptake assay (Ziambaras et al., 1998). In this assay, cells labeled with both a gap junction-permeable dye (calcein red/orange AM, shown in black) and a non-spreading membrane dye (DiD, shown in magenta) were added to a subconfluent monolayer of unlabeled cells. The formation of GJs is indicated by membrane dye-negative cells that become calcein positive with time. HeLa cells express low levels of endogenous connexins (Elfgang et al., 1995) and display minimal dye coupling (**Fig. 2B**). However, stable expression of Cx46 or transient expression of Cx43 in HeLa cells established functional gap junctions and coupling between cells, as evidenced by the spread of calcein dye (shown in black) between cells (**Fig. 2B**). Using stable Cx46-expressing HeLa cells, we then screened the 727 compounds of the NIH Clinical Collection of FDA-approved small molecules for their ability to inhibit Cx46-mediated GJIC at a concentration of 10 µM over a treatment time of 3 hours (**Fig. 2C**). The spread of calcein between treated cells was imaged after 5 hours and compared to both vehicle (DMSO) treatment and treatment with the pan-gap junction inhibitor carbenoxolone (CBX; 200 nM). We identified a number of compounds that blocked Cx46-mediated GJIC compared to carbenoxolone as a positive control (**Fig. 2D**). Several of the top hits were further screened at concentrations between 0.1 µM and 10 µM found that the FDA-approved anti-mycobacterial drug clofazimine inhibited Cx46 GJIC at the lowest concentrations compared to the other hits (**Fig. 2E**), with little effect on Cx46 hemichannel activity (**Fig. 2F**). Together, these results demonstrate that clofazimine is a candidate to inhibit Cx46 GJIC without affecting potential hemichannel activity.

**Figure 2.**
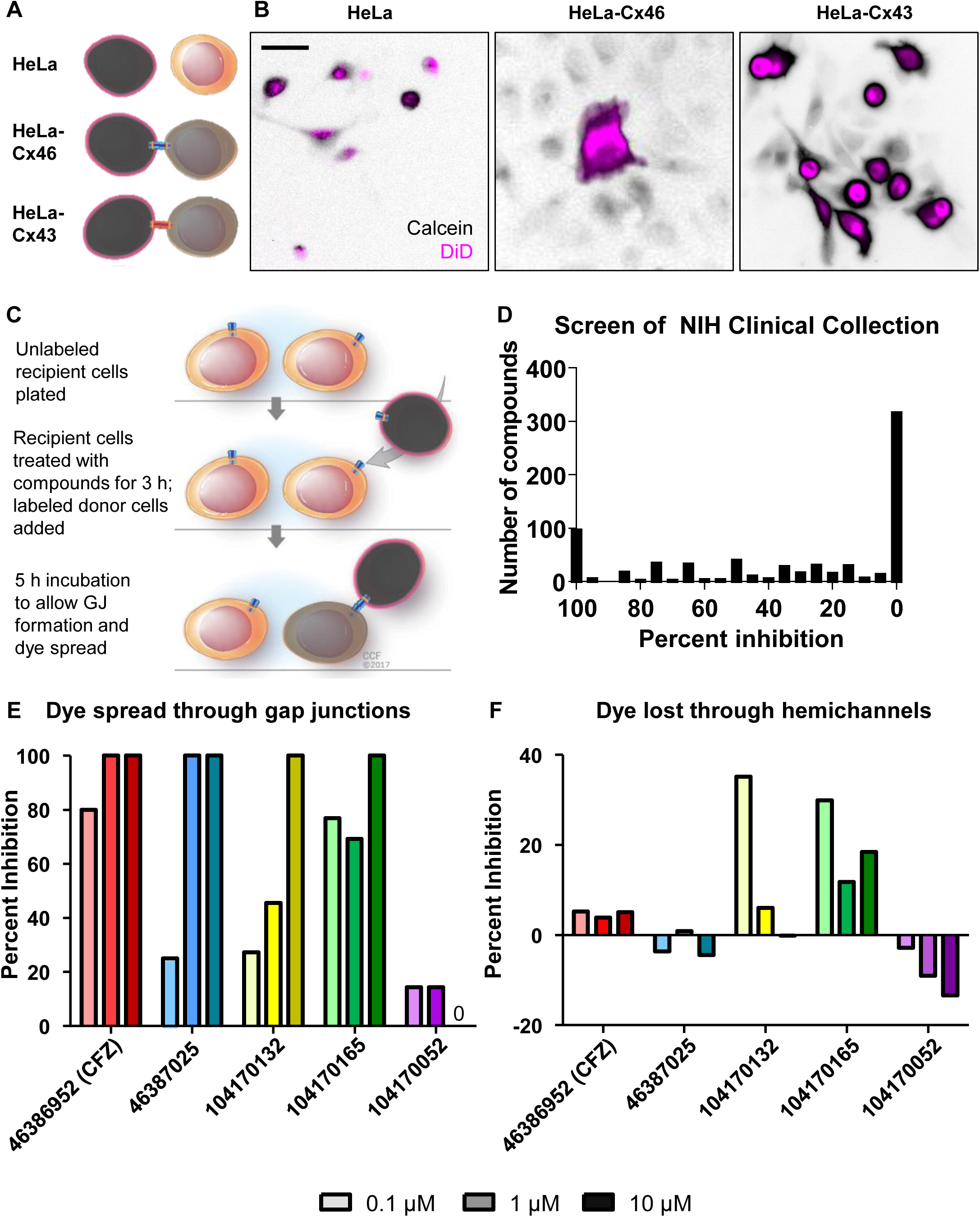
A screen of FDA-approved small molecules identifies clofazimine as an inhibitor of Cx46-mediated cell-cell communication. (**A**) Schematic of calcein dye transfer between HeLa cells expressing no exogenous connexin proteins and HeLa cells transfected with Cx43 or Cx46. Cells are labeled with Vybrant DiD (pseudocolored magenta), which cannot pass between cells, and calcein red/orange AM (pseudocolored black), which spreads between cells through gap junctions. (**B**) Parachute dye transfer assay of parental HeLa cells, stable Cx46-expressing HeLa cells, and transiently transfected Cx43-expressing HeLa cells. Unlabeled cells were plated in a subconfluent monolayer, and dual-labeled cells were added. If gap junctions formed between labeled and unlabeled cells, the calcein dye (shown in black) diffused into cells that were not labeled with DiD (magenta). “0” on the plot indicates no inhibition was observed. Scale bar, 50 µm. (**C**) Schematic of the parachute dye transfer assay with timing used to test the NIH Clinical Collection compounds for inhibition of Cx46-mediated cell-cell communication in stable HeLa-Cx46 cells. A subconfluent monolayer of HeLa-Cx46 cells was plated and incubated with drugs at 10 µM for 3 h. A separate population was labeled with calcein red-orange AM and Vybrant DiD and added to the recipients, and dye transfer was imaged after 5 h. (**D**) Summary graph of the degree to which the drugs from the NIH Clinical Collection inhibited Cx46-mediated GJIC. Percent inhibition is relative to DMSO vehicle control treatment (0%) and the pan-gap junction inhibitor carbenoxolone (100%). (**E-F**) Validation of the screen results. Cells were treated with increasing concentrations (0.1 µM, 1 µM, 10 µM) of four top hits from the screen and one hit that did not show inhibition (purple). Those cells were then either plated and incubated with a labeled population of donor cells (**E**) to measure GJIC or plated sparsely (**F**) to assay dye leakage through hemichannels. Data are normalized to DMSO (0% inhibition) and carbenoxolone (100%), and these experiments were performed in triplicate.

### Cx46 is more sensitive than other connexins expressed in GBM to inhibition by clofazimine

To specifically target Cx46 in CSCs, the lead compound should have limited efficacy against the other 20 human connexins. To test the specificity of clofazimine for Cx46, we first screened for the additional connexins expressed in GBM using bioinformatics. Using both RNA-sequencing and microarray data from the GlioVis database (http://gliovis.bioinfo.cnio.es/), we identified the connexins most highly expressed in GBM compared to normal brain (**Fig. 3A**). In addition to Cx46, which was the most highly expressed relative to normal brain tissue, Cx45 and Cx37 were also detected at higher levels in GBM. We also screened clofazimine against Cx43, the most ubiquitously expressed connexin throughout the body (Oyamada et al., 2005). HeLa cells expressing any of these four connexins displayed GJ coupling, as evidenced by the spread of calcein dye (black) from DiD (shown in magenta)-labeled donor cells to unlabeled recipient cells (**Fig. 3B**). As expected, the pan-gap junction inhibitor CBX inhibited calcein spread for each connexin. However, while coupling of HeLa cells expressing Cx46 was blocked by clofazimine, cells expressing Cx43, Cx37, and Cx45 continued to exhibit GJIC even in the presence of clofazimine (**Fig. 3C**). These data indicate that of the connexins tested, clofazimine was specific for inhibition of Cx46-mediated GJIC.

**Figure 3.**
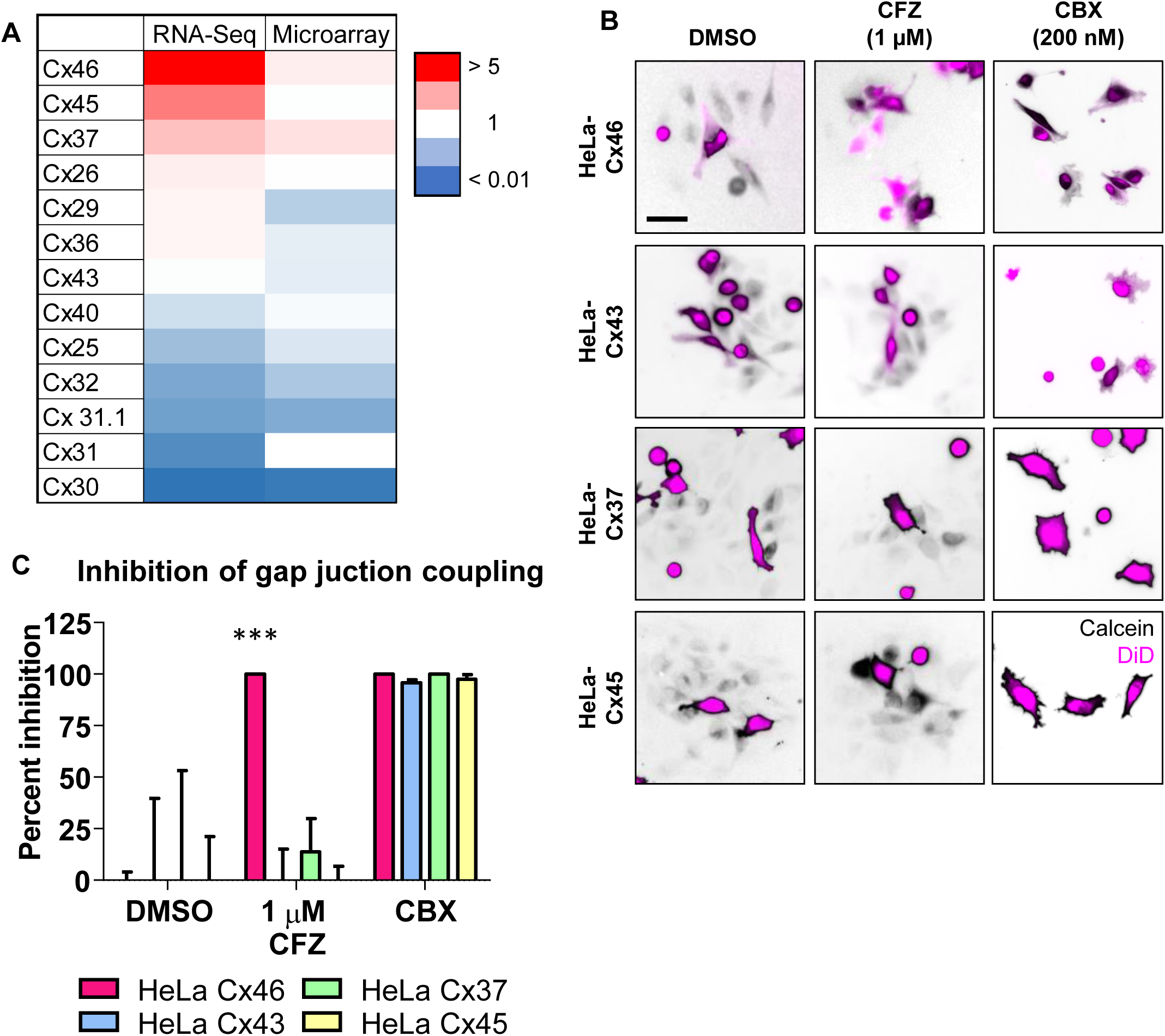
Cx46 is more sensitive than other connexins expressed in GBM to inhibition by clofazimine. (**A**) Heatmap of connexin mRNA expression in GBM compared to normal brain tissue by both RNA-sequencing and microarray. Data are from TCGA and were obtained from the GlioVis. Red indicates higher expression compared to normal brain, while blue indicates lower expression than normal brain tissue. (**B**) Parachute dye transfer assay of HeLa cells expressing different connexin proteins. HeLa cells were transfected with connexin proteins, unlabeled cells were plated in a subconfluent monolayer, and cells dual labeled with Vybrant DiD (shown in magenta) and calcein red/orange AM (shown in black) were treated with DMSO, 1 µM clofazimine (CFZ), or 200 nM carbenoxolone (CBX) for 3 h and added to the unlabeled cells. The presence of calcein dye (black) in cells that are not magenta indicates GJIC. Scale bar, 50 µm. (**C**) Quantification of B. The percent inhibition of GJIC with clofazimine is shown compared to that of vehicle and the pan-gap junction inhibitor carbenoxolone. *** p<0.001 by unpaired Student’s t-test with Welch’s correction compared to the DMSO-treated control. Data are represented as mean ± SEM. n = 3.

### Clofazimine preferentially targets GBM CSCs compared to non-CSCs

Based on our previous studies identifying Cx46 as an essential connexin expressed by GBM CSCs and our above results that clofazimine preferentially inhibits coupling of cells expressing Cx46, we hypothesized that clofazimine would specifically target GBM CSCs compared to non-CSCs. Treatment of CSCs and non-CSCs with increasing concentrations of clofazimine from 0.05 µM to 5 µM allowed us to calculate IC_50_ values of approximately 2 µM for the CSC population of four different patient-derived xenograft specimens (**Fig. 4A**). In contrast, the non-CSC population never reached 50% growth inhibition within the same concentration range of clofazimine. For comparison, the IC_50_ of the immortalized, non-transformed fibroblast cell line NIH3T3 was measured at approximately 86 µM, indicating that CSC growth was dramatically more sensitive than that of other cell types to clofazimine. Limiting dilution analysis showed a significant and striking effect of clofazimine on CSC self-renewal, even at concentrations where proliferation was only minimally affected (0.5 µM; **Fig. 4B and Supplemental Fig. S2A**). This inhibition of CSC growth and self-renewal was accompanied by a concentration-dependent increase in apoptosis in the CSC population, with minimal induction of apoptosis in the non-stem cells (**Fig. 4C**).

**Figure 4.**
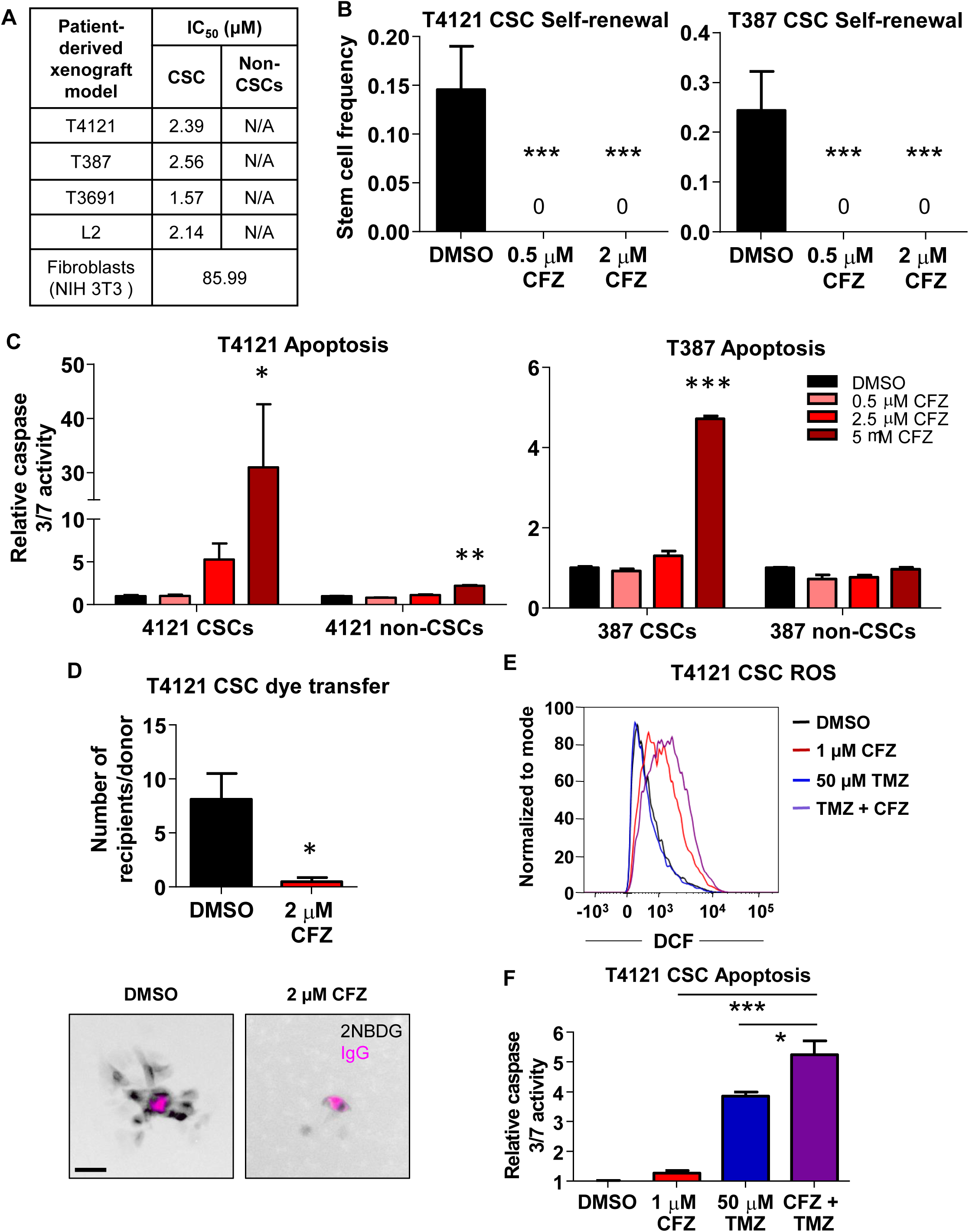
Clofazimine preferentially targets GBM CSCs compared to non-CSCs. (**A**) Summary of IC_50_ values for clofazimine (CFZ) in four different patient-derived xenograft matched CSCs and non-CSCs and the NIH3T3 untransformed fibroblast cell line. Cells were treated with increasing concentrations of clofazimine for 3 d, and cell number was measured using CellTiter-Glo. Because non-CSCs never reached a 50% decrease in cell number, an IC_50_ value could not be calculated. n = at least 3 experiments with cells plated in triplicate. Data are represented as mean ± SEM. n = 3. (**B**) CSCs from two different patient-derived xenograft specimens were plated into drug-containing medium at increasing cell densities (1, 5, 10, 20 cells/well of a 96-well plate), and the number of wells containing spheres was counted after 10-14 days. The online algorithm described in the Methods section was used to calculate stem cell frequency. *** p<0.001 by χ^2^ test compared to the DMSO-treated control. Data are represented as mean ± range. n = 3 experiments, with 24 technical replicates per cell number per experiment. (**C**) CSCs and non-CSCs from two different patient-derived xenograft specimens were treated with clofazimine for 3 d, and active caspase 3/7 was measured using Caspase-Glo. The values shown are normalized to the number of total cells at the same time point and are relative to the DMSO control for each cell type. * p<0.05, ** p<0.01, *** p<0.001 by unpaired Student’s t-test with Welch’s correction compared to the respective DMSO-treated control. Data are represented as mean ± SEM. n = 2 experiments, each performed in triplicate. (**D**) CSCs were plated in a subconfluent monolayer on Geltrex, treated for 24 h, and microinjected with 2-NBDG (pseudocolored black) and a far-red fluorescently labeled IgG (pseudocolored magenta). Cells were imaged over 2 h, and the number of cells receiving 2-NBDG from each donor cell was quantified. * p<0.05 by unpaired Student’s t-test with Welch’s correction compared to the DMSO-treated control. Data are represented as mean ± SEM. n = 8 donors over 7 fields (DMSO) and n = 4 donors over two fields (clofazimine). Scale bar, 50 µm. (**E**) Flow cytometry was used to measure the amount of fluorescent DCF produced from H_2_CDFDA as a measurement of ROS produced. CSCs were treated concurrently for 24 h with 50 µM temozolomide (TMZ) and for 16 h with 1 µM clofazimine, manually removed from the plate using a cell scraper, and subjected for flow cytometry. Representative data from one of n = 3 experiments are shown. (**F**) Cells were plated in 96-well plates and treated as in E. Active caspase 3/7 was measured using Caspase-Glo. Data are normalized to the total number of cells at that time and are shown relative to the DMSO-treated control. * p<0.05, *** p<0.001 by unpaired Student’s t-test with Welch’s correction compared to treatment with clofazimine alone. Data are represented as mean ± SEM. n = 2 experiments, each performed in triplicate. See also Supplemental Figure S2.

Based on our data that clofazimine inhibited dye coupling in HeLa cells expressing Cx46 and not other connexins (**Fig. 3B**), we hypothesized that clofazimine was similarly acting through an inhibition of GJIC in CSCs. Indeed, treatment with clofazimine inhibited the spread of the fluorescent glucose analog 2-NBDG microinjected in CSCs compared to vehicle (**Fig. 4D**), confirming that clofazimine is able to inhibit GJIC in CSCs. To further test whether clofazimine induced additional off-target effects, we performed RNA-sequencing on CSCs from xenograft specimen T4121 treated with 2 µM clofazimine for a short time period of 6 hours. Increases and decreases in transcript expression with treatment compared to vehicle were relatively modest, with changes falling within 3-fold of the value of the vehicle-treated samples (**Supplemental Fig. S2B-C**). We performed functional gene annotation and pathway enrichment analysis on the top differentially expressed genes (https://david.ncifcrf.gov/) and found no significant pathway enrichment within reported gene groups with clofazimine treatment, suggesting limited off-target effects with clofazimine treatment. Clofazimine has also been reported to target GBM cells by affecting the function of the membrane potassium channel Kv1.3, which is highly expressed in many cancer cell lines compared to normal tissue (Leanza et al., 2015; Venturini et al., 2017). We therefore tested our CSCs and non-CSCs to determine whether higher levels of Kv1.3 in the CSCs could be responsible for their sensitivity to clofazimine. However, GBM CSCs from the patient-derived xenograft T4121, which are more sensitive to clofazimine than their non-stem counterparts, expressed approximately 4-fold less Kv1.3 transcript than non-CSCs (**Supplemental Fig. 2D**), suggesting that the enhanced sensitivity to clofazimine of CSCs is likely not due to Kv1.3 channels.

Inhibition of GJs has been reported to increase the cellular levels of reactive oxygen species (ROS) (Giardina et al., 2007; Le et al., 2014; Zundorf et al., 2007). As expected, treatment with 1 µM clofazimine for 3 days led to an increase in intracellular ROS as measured by production of fluorescent 2’,7’-dichlorofluorescein (DCF) from 2’,7’-dichlorodihydrofluorescein diacetate (H_2_DCFDA) and detected using flow cytometry (**Fig. 4E**). Based on our observations that clofazimine is toxic to GBM CSCs, we combined clofazimine with temozolomide, GBM standard-of-care chemotherapy. Temozolomide alone (50 µM) did not increase ROS compared to DMSO vehicle treatment, but a combination of temozolomide with clofazimine further increased ROS above the level observed for clofazimine alone. This increase in ROS was accompanied by a significant increase in apoptosis in cells treated with both temozolomide and clofazimine compared to either compound alone (**Fig. 4F**), and this increase in the combination treatment was greater than an additive effect, suggesting that clofazimine sensitizes CSCs to chemotherapy. Together, these results indicate that clofazimine inhibits GBM CSC growth, survival, and self-renewal, likely through its effects on Cx46-mediated GJIC, and combines with GBM standard-of-care therapies to further increase tumor cell death.

### Clofazimine decreases tumor growth in vivo

The current World Health Organization (WHO) dosing schedule of clofazimine for multibacillary leprosy includes one monthly dose of 300 mg and an additional 50 mg daily in combination with the drugs dapsone and rifampicin for a period of 12 months (Fischer, 2017). To determine whether clofazimine inhibits tumor growth in vivo, we selected a dosage equivalent to the maximum recommended daily human dose (Novartis, https://www.drugs.com/pro/lamprene.html), 200 mg/day (2.44 mg/kg based on an average human body weight of 80 kg), solubilized in corn oil and delivered via intraperitoneal injection (IP). At this dose, we were unable to detect significant levels of clofazimine in treated animals compared to the background levels observed in the brain of vehicle-treated animals (**Fig. 5A** and **Supplemental Fig. S3A**), and we also observed low penetration of the blood-brain barrier by clofazimine in mice (**Fig. S3B**). For these reasons, rather than treating mice with intracranial tumors, we instead treated animals bearing flank tumors generated by implantation of CSCs from the PDX specimen T4121. Clofazimine administration began once all animals presented palpable tumors. Treatment with 2.44 mg/kg clofazimine by IP for two weeks led to a significant decrease in tumor growth as measured using digital calipers (**Fig. 5B**), with the final tumor size shown in **Fig. 5C**. As the normal tissue distribution of Cx46 is primarily in the lens, we also tested whether inhibition of Cx46 had an effect on animal vision and observed no significant changes compared to treatment with vehicle (**Supplemental Fig. S3C**). Together, these results indicate that clofazimine targeting of Cx46-mediated GJIC is able to slow tumor growth without impacting other major Cx46 functions, including vision.

**Figure 5.**
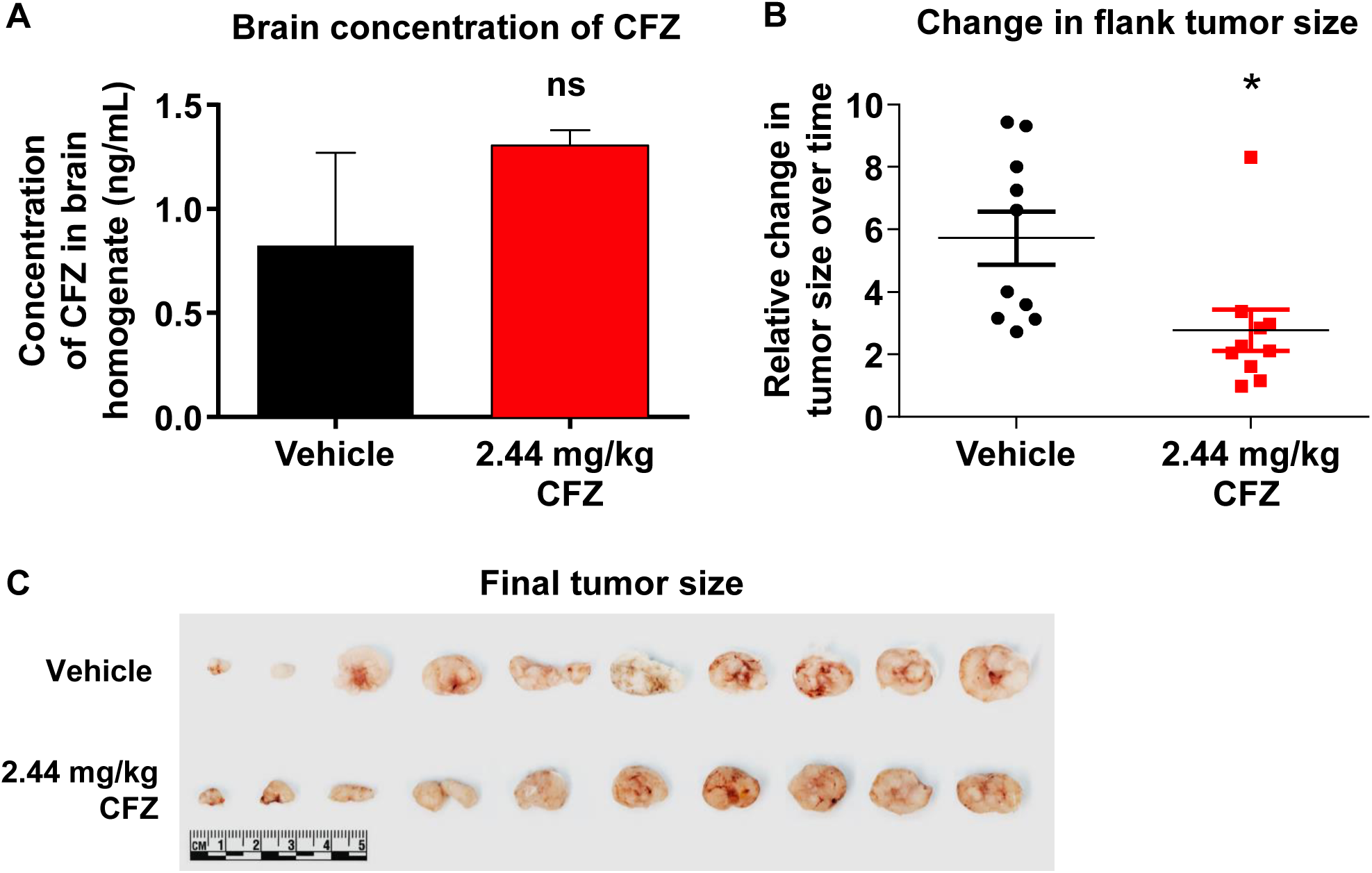
Clofazimine decreases tumor growth in vivo. (A) Female NSG mice were treated with 2.44 mg/kg clofazimine (CFZ) in 200 µl corn oil by intraperitoneal (IP) injection daily for 5 d. On day 8, animals were euthanized, and brains were homogenized in PBS and subjected to mass spectrometry for clofazimine. n = 3 brains each for vehicle and clofazimine, and data are shown as mean ± SEM. The difference is not significant (ns) by unpaired Student’s t-test with Welch’s correction compared to treatment with vehicle alone (background signal). (**B**) Female NSG mice were injected with 1×10^6^ T4121 CSCs into their right flanks. Three weeks later, when tumors became palpable, animals were treated with 2.44 mg/kg clofazimine in corn oil IP for two weeks. Tumor size was measured using digital calipers, and the change over time is given as the measurement on day 12 relative to day 0. Tumors were excised on day 15, and the final tumor sizes are shown in (**C**). The scale is shown in the figure. * p<0.05 by unpaired Student’s t-test with Welch’s correction compared to treatment with vehicle (corn oil). n = 10 mice per arm. All data points as well as the mean and SEM are shown. See also Supplemental Figure S3.

## Discussion

Connexin proteins serve three main cellular functions: exchange of small molecules between cells, exchange of small molecules between cells and the extracellular space, and mediating intracellular protein-protein interactions. We previously showed that Cx46 is required for GBM CSC proliferative ability, survival, self-renewal, and tumor formation (Hitomi et al., 2015). Here, using point mutations that disrupt specific functions of the protein, we show that the essential function of Cx46 in these cells is the formation of functional Cx46 GJs. It remains an open question as to the key tumor cell and CSC mediators that pass through GJs, which likely include a combination of ions (K^+^, Ca^2+^, Na^+^), ROS and antioxidants, metabolites such as glucose, cAMP, and non-coding and microRNAs (Lim et al., 2011; Loewenstein and Kanno, 1964; Patel et al., 2016). Our results contrast with the hypothesis that aberrant hemichannel activity of connexins underlies their role in pathologies (Kim et al., 2016; Leybaert et al., 2017) and suggest that therapies designed to target GJIC mediated by specific connexins may be valuable for certain diseases, including GBM.

To this end, we screened FDA-approved compounds for Cx46 GJIC inhibitors and identified the anti-leprosy drug clofazimine, which inhibited GBM CSC cell-cell communication; decreased CSC growth, survival, and self-renewal; and decreased tumor growth in a subcutaneous tumor model. Although pan-gap junction inhibitors are available clinically and have shown efficacy in our models (Hitomi et al., 2015), specific inhibitors for connexin isoforms have yet to be identified or developed. The majority of connexin modulators developed so far, most of which are designed to target Cx43 or multiple connexin isoforms, are peptide mimetics that interrupt a specific binding activity of the molecule – either within the molecule or between molecules – and thus affect protein or channel function (Jaraiz-Rodriguez et al., 2017; Naus and Giaume, 2016). Although little is known about precisely how these mimetics modulate connexin activity, they possess varying efficiencies at inhibiting and/or stimulating both GJ and hemichannel activity (Evans et al., 2012; Wang et al., 2013). However, due to the homology among connexin isoforms, many of these mimetics fail to exhibit specificity for a specific connexin. In contrast, we show that the small molecule clofazimine is specific for Cx46 compared to Cx43, Cx45, and Cx37. Few small molecules have been identified to target connexins; those that have been developed increase GJIC in astrocytes or specifically target hemichannels, neither of which are relevant to blocking Cx46-mediated GJIC in GBM CSCs (Naus and Giaume, 2016).

Previous studies described an inhibitory role for clofazimine in GBM cells. Significant apoptosis has been observed in conventional GBM cell lines treated with clofazimine, and this cell death was attributed to inhibition of the mitochondrial membrane ion channel Kv1.3 (Venturini et al., 2017). We observed similar cell death of GBM CSCs upon treatment with clofazimine, with little effect on non-CSCs. However, we also detected 4-fold higher levels of Kv1.3 transcript in the clofazimine-resistant non-CSC population, suggesting that clofazimine does not act through Kv1.3 inhibition in our hands. Clofazimine was also previously identified in a screen to inhibit growth of the conventional GBM cell line U87 (Jiang et al., 2014). In contrast, rather than screening for compounds that inhibit GBM cell growth in culture, we identified a CSC essential process, Cx46-mediated GJIC, and screened for inhibitors of this mechanism. Our future work will investigate the mechanism by which clofazimine blocks Cx46-mediated intercellular communication. Based on our observations that the cysless mutant inhibits CSC maintenance similarly to clofazimine and that few transcripts were altered by short-term treatment, we speculate that the drug could act extracellularly to physically block the channel opening or hemichannel-hemichannel docking. However, it remains possible that clofazimine functions in another manner, for example by altering membrane permeability, mitochondrial function, or cell signaling.

Although clofazimine shows promise for treating GBM, there are several challenges to its therapeutic use. We show that clofazimine exhibits minimal penetration of the blood-brain barrier, and its low solubility and high lipophilicity are also barriers to translation for brain tumors. There has been conflicting evidence for whether clofazimine is able to penetrate the brain; while some studies have reported no detectable levels in the brain (Baik et al., 2013; Holdiness, 1989), other studies detected a level of 156 ng/mL of clofazimine in the brain of mice treated with 25 mg/kg of the drug (Baijnath et al., 2015) and an effect on Kv1.3 channels in the brain in animals treated with 50 mg/kg clofazimine after traumatic brain injury (Reeves et al., 2016). In contrast, using the equivalent of the maximum tolerated human dose (2.44 mg/kg in mice), we were unable to detect clofazimine in the brain using mass spectrometry or the ability for it to cross the blood-brain barrier. This difference may be due to differences in delivery route, solvent, or concentration. In a previous report, clofazimine failed to inhibit growth of intracranial syngeneic mouse gliomas (Venturini et al., 2017), which is supported by our observations that clofazimine at human-relevant doses does not cross the blood-brain barrier. Medicinal chemistry derivatization of clofazimine to optimize solubility and blood-brain barrier penetration will allow us to develop a more optimal analog of clofazimine for further pre-clinical and clinical testing and could lead to improved next-generation therapies with reduced side effects for patients with GBM.

## Experimental Procedures

### Origin of cells

Established GBM xenografts T4121, T3691, and T387 were previously reported (Alvarado et al., 2016; Bao et al., 2006; Schonberg et al., 2015) and were obtained via a material transfer agreement from Duke University. L2 cells were obtained from the University of Florida (Deleyrolle et al., 2011; Siebzehnrubl et al., 2013). All human GBM samples were originally established under an IRB-approved protocol that facilitated the generation of xenografts in a de-identified manner from excess tissue taken from consented patients. GBM cells were passaged in immune-deficient NOD.Cg-Prkdc^scid^Il2rg^tm1Wjl^/SzJ (NSG) mice (obtained from The Jackson Laboratory, Bar Harbor, ME, USA) and dissociated from established mouse xenografts under Cleveland Clinic-approved protocols. Six-week-old female mice were unilaterally injected subcutaneously in the flank with freshly dissociated human GBM cells, and animals were sacrificed by CO_2_ asphyxiation and secondary cervical dislocation when tumor volume exceeded 5% of the animal’s body weight. HeLa and NIH3T3 cells were obtained from ATCC.

### Cell culture

Xenograft tumors were dissociated using papain (Worthington Biochemical Corporation, Lakewood, NJ) and cultured overnight in supplemented neurobasal medium (neurobasal medium (Life Technologies) with 2% B27 (Life Technologies), 1% penicillin/streptomycin (Life Technologies), 1 mM sodium pyruvate (Life Technologies), 2 mM L-glutamine, 20 ng/mL EGF (R&D Systems, Minneapolis, MN, USA), and 20 ng/mL FGF-2 (R&D Systems)). T4121, T3691, and T398 xenografts were sorted for CD133+ and CD133-populations using the CD133 Magnetic Bead Kit for Hematopoietic Cells (CD133/2; Miltenyi Biotech, San Diego, CA, USA). CD133+ cells were maintained in supplemented neurobasal. CD133-cells were maintained in DMEM with 5% FBS and 1% pen/strep. L2 cells were maintained in these divergent media conditions without sorting.

HeLa and NIH3T3 cells were maintained in DMEM with 10% FBS and 1% pen/strep. The HeLa-Cx46 stable cell line was cultured with the addition of 400 µg/mL G418. All cells were grown in a humidified incubator at 37°C with 5% CO_2_.

### Plasmids and DNA constructs

The Cx46 expression vector was created by inserting the Cx46 cDNA (catalog# RDC0535, R&D Systems) between the HindIII and XbaI sites of pEGFP-N3, excising the GFP tag. This backbone was used for site-directed mutagenesis to introduce the L11S, T19M, and cysless mutations, using the primers shown in Table 1.

The primers for cysless were designed so that the PCR reactions must be performed sequentially from N-terminus to C-terminus.

pLPCX-Cx43-IRES-GFP was obtained from Addgene (#65433). pcDNA3.1/Hygro(+)-GJC1 (Cx45; cloneID: OHu04829) and pcDNA3.1/Hygro(+)-GJA4 (Cx37; cloneID: OHu33346) were obtained from GenScript.

### Transfection and establishment of HeLa-Cx46 stable cell line

For GBM CSC transfections, 1×10^6^ cells were plated per well of a 6-well plate adherently on Geltrex (Thermo Fisher Scientific) to obtain a confluence of approximately 75-80%. Six hours later, cells were transfected with Cx46 or its mutant forms using FuGENE HD (Promega) according to the manufacturer’s protocol. Briefly, cells were transfected with 5 µg total DNA (4 µg of connexin and 1 µg pEGFP-N3 to track transfection efficiency) using 15 µl FuGENE per well. The following day, cells were removed from the plate using Accutase (BioLegend) and plated for downstream assays. pEGFP-N3 was used as a vector control.

HeLa cells were seeded at 400,000 cells in a 6-well plate and transfected using XtremeGene HP (Roche) according to the manufacturer’s protocol. In brief, each well received 2 ug of DNA and 6 uL of XtremeGene reagent. Dye-transfer recipients were plated 24 hours after transfection, and donors were plated and images taken at 48 hours post-transfection. Stable HeLa-Cx46 cells were derived by transfecting HeLa cells with Cx46 (without the GFP tag). Cells were selected with G418 (400 µg/mL), and single-cell clones were tested for the ability to exhibit dye coupling.

### Compounds

Clofazimine was obtained from Sigma-Aldrich (catalog # C8895) and solubilized in DMSO at a concentration of 10 mM for in vitro experiments and at 0.489 mg/mL in corn oil for in vivo experiments.

### Proliferation and apoptosis

For proliferation, IC_50_, and apoptosis assays, 2,000 cells were plated per well of a white-walled 96-well plate in triplicate. The number of cells was measured using CellTiter-Glo (Promega) on days 0, 1, 3, 7, and 10 according to the manufacturer’s protocol using ATP content as a surrogate of cell number, and apoptosis was measured using CaspaseGlo 3/7 (Promega) on days 1 and 3 according to the manufacturer’s protocol. For the proliferation of GBM CSCs in the presence of Cx46 and Cx46 mutants, similar results were obtained using the DNA-based CyQUANT Direct Cell Proliferation Assay Kit (Thermo Fisher Scientific). For drug treatments, cells were seeded in triplicate at 2,000 cells per well of a 96-well plate, and the appropriate concentration of drug was added 6-24 hours later. Cells were analyzed both at 0 and 72 h after treatment with drug.

### Limiting dilution analysis

CSCs were dissociated using Accutase and plated in a 96-well plate at increasing cell numbers (1, 5, 10, and 20 cells/well) with 24 replicates per cell number. Cells were plated into drug-containing media, and the number of wells containing spheres was counted after 10-14 days. An online algorithm (http://bioinf.wehi.edu.au/software/elda/) (Hu and Smyth, 2009) was used to calculate stem cell frequency. This was repeated at least two times per drug concentration. cDNA and qPCR

For qPCR, RNA was extracted from cells using TriZOL (Life Technologies) according to the manufacturer’s protocol. A total of 1 µg of RNA was used for reverse transcription using a qScript cDNA Synthesis Kit (QuantaBio) according to the manufacturer’s recommendations. Equal volumes of cDNA were amplified using Fast SYBR® Green Master Mix (Applied Biosystems) on a Step-One Plus Real-Time PCR system (Applied Biosystems). Data were analyzed using the ΔΔCt method to calculate relative levels of product. qPCR primers are provided in Supplemental Table 1.

### Screen for Cx46 inhibitors in the NIH Clinical Collection

Non-labeled Cx46-HeLa cells were seeded at 20,000 cells per well in a 96-well plate in DMEM with 10% FBS and 1% pen/strep. The following morning, drugs were added to a concentration of 10 µM to 80 of the wells, leaving 16 for positive and negative inhibition controls. Carbenoxelone (200 nM) was used a positive control for dye transfer inhibition, while negative control wells were left untreated. Separately, a population of calcein red-orange AM/Vybrant DiD dual-labeled Cx46-HeLa cells was generated. These cells were incubated in serum-free DMEM containing calcein red-orange (resuspended in 50 µL of DMSO and used at 1:1000; Thermo Fisher Scientific) and Vybrant DiD (1:500, Thermo Fisher Scientific) at 37°C for 1 h. Following a 3 h incubation of the unlabeled recipients with drug, the dual-labeled donor population was added at a concentration of 3,000 cells/well. These cells were incubated together at 37°C for 5 h and then imaged after 5 hours. Each plate contained 80 drugs and 16 controls, accounting for 9 experimental runs. Each drug was screened one time per drug as a cursory screen. Following the identification of possible targets, a secondary screen of a selection of top hits that were visually verified and readily available was performed at drug concentrations of 10 µM, 1 µM, and 0.1 µM.

For screen quantification, calcein fluorescence (red) was used to create a mask to eliminate any cells left entirely unlabeled and any background fluorescence. The Vybrant DiD fluorescence (far red) image was used to create another binary mask to define DiD-positive donor cells. These mask images were given values of 0 (no dye present) or 1 (dye present) and then multiplied by the calcein image. ImageJ particle analysis of the resulting product images provided us with the raw integrated density (RID) of the total calcein dye per imaged cell. The sum of the particle analysis of the product of the calcein mask and the calcein image gave the total calcein amount, and that of the product of the DiD mask and the calcein image gave the amount of calcein retained in the donor cells. Percent transfer was calculated by (total calcein – retained calcein)*100/total calcein.

For hemichannel function assessment, labeled populations were generated as described above and seeded at 3,000 cells per well. Cells were given an hour to adhere and then imaged every 15 minutes for 5 hours. Loss of calcein through hemichannels was quantified as the percent of dye that was lost at after 5 h compared to time 0.

For HeLa cells expressing different connexin proteins, cells were prepared and imaged as stated above. Images were quantified as the number of unlabeled cells (recipients) receiving calcein dye per donor cell.

For microinjection of CSCs, subconfluent monolayers of cells plated on Geltrex-coated glass coverslips in 35 mm dishes were pretreated for 16 h with the indicated concentration of clofazimine in growth media. Cells were then injected with far-red fluorescent IgG and the fluorescent glucose analog 2-(N-(7-nitrobenz-2-oxa-1,3-diazol-4-yl)amino)-2-deoxyglucose (2-NBDG) as described (Hitomi et al., 2015) and imaged as above. Images were again quantified as the number of unlabeled cells (recipients) receiving calcein dye per donor cell.

### GlioVis analysis of connexins in GBM

The Cancer Genome Atlas (TCGA) dataset was interrogated using GlioVis (gliovis.bioinfo.cnio.es, citation) for microarray (Agilent-4502A) and RNAseq levels of all available connexin genes. Relative levels of non-tumor and GBM tissues were analyzed, and the fold change is represented as a heat map.

### RNA sequencing

T4121 CSCs were treated with clofazimine at 2 µM for 6 hours and lysed for RNA using a Macherey-Nagel Nucleospin RNA isolation kit. RNA-seq libraries were prepared using ∼10,000 ng of total RNA. Briefly, the protocol included PolyA+ RNA selection, cDNA synthesis, end repair, A-base addition, and ligation of the Illumina-indexed adapters according to previously published methods (Zhang et al., 2012). Total transcriptome libraries were prepared as previously described. Library quality and quantity were measured on an Agilent 2100 Bioanalyzer for product size and concentration. Libraries were also precisely quantified by using a KAPA Library Quantification kit prior to loading on the sequencer and pooled at equimolar quantities between samples. Single-end libraries were sequenced with the Illumina HiSeq 2500 (1×5 read length), with sequence coverage up to 20 M total reads.

Single-end transcriptome sequencing reads were aligned to the human reference genome (GRCh37/hg19) using the spliced read mapper TopHat2 (TopHat 2.0.4) (Kim et al., 2013). Gene expression, as fragments per kilobase of exon per million fragments mapped (FPKM; normalized measure of gene expression), was calculated using Cufflinks (Trapnell et al., 2012). We considered differential expression of the gene when the calculated p<0.01 and there was a 1.5-fold difference (increase or decrease).

The database for annotation, visualization and integrated discovery (DAVID) analysis was used for functional clustering and annotation of differentially expressed genes (http://david.abcc.ncifcrf.gov/) (Jiao et al., 2012). DAVID is a web-based online bioinformatics resource that aims to provide tools for pathway mining and the subsequent functional interpretation of large lists of genes/proteins using a comprehensive and exhaustive set of knowledge-base libraries. The publication on the DAVID webserver suggests investigating clusters with an enrichment score ≥ 1.3, while our highest enrichment score was 1.06, suggesting no major disturbance of any functional pathway/gene ontology group.

### Reactive oxygen species (ROS)

To measure intracellular ROS, CSCs were treated with 50 µM temozolomide for 24 h and 1 µM clofazimine for 16 h. Cells were then collected and incubated with 1 µM H_2_DCFDA (Life Technologies) for 15 min at 37°C. Cells were then washed twice in PBS, and the green fluorescent DCF produced was analyzed on a BDFortessa flow cytometer. DAPI exclusion was used to gate for live cells, and H_2_O_2_ was used as a positive control for ROS production.

### Orthotopic tumors

Six-to eight-week-old immunocompromised female NSG mice were injected with 1×10^6^ CSCs from the patient-derived xenograft T4121 into their right flank. Three weeks later, when tumors were palpable, mice were treated IP with clofazimine at 2.44 mg/kg in corn oil or vehicle for two weeks on weekdays. Tumor width was measured using digital calipers on days 1 and 12 and is provided as the relative change over that time. Animals were sacrificed on day 15, and tumors were excised and imaged. All animal experiments were performed under Cleveland Clinic-approved Institutional Animal Care and Use Committee-approved protocols.

### Blood-brain barrier

To assess the permeation of clofazimine into normal brain tissue, mice were intraperitoneally injected with 100 µL of a 25 mg/mL suspension of clofazimine in corn oil or vehicle. After 10 minutes of circulation, mice were euthanized, and brains were extracted, snap frozen in isopentane, and sliced into 20 µm sections. Slides were analyzed using a MVX10 MacroView microscope (Olympus) equipped with an ORCA_Flash4.0 v2 sCMOS fluorescent camera (Hamamatsu). A linear range of standards in the brain was developed with varying concentrations (16 µg/mg to 2.5 mg/mg).

### Mass spectrometry

Brains from mice treated IP with 2.44 mg/kg clofazimine were excised and homogenized in PBS. For mass spectrometry, clofazimine was used as the internal standard. Brain homogenate (50 µL) was mixed with 150 µL methanol and then centrifuged at 12,000 x g for 10 min. The supernatant (100 µL) was transferred to an HPLC vial. For LC/MS/MS analysis of clofazimine, 2 µL supernatant was injected into a Shimadzu LCMS-8050 for quantitation of clofazimine. A gradient with a flow rate of 0.3 mL/min was used to separate clofazimine by reverse-phase chromatography using a Prodigy C18 column (2.1 × 50 mm, 5 µm) from Phenomenex. The mobile phases were A (water containing 5 mM ammonium acetate) and B (methanol containing 5 mM ammonium acetate). The run started with 70% mobile phase B from 0 to 2 min. Solvent B was then increased linearly to 100% B from 2 to 6 min and held at 100% B from 6 to 12 min. The column was finally re-equilibrated with 70% B for 7 min. The HPLC eluent was directly injected into a triple quadrupole mass spectrometer (Shimadzu LCMS-8050), and the clofazimine was ionized at ESI positive mode, using selected Reaction monitoring (SRM). The SRM transitions (m/z) were 474 to 432. For data analysis, the software Labsolutions was used to process the data and obtain the peak areas for clofazimine. The external standard calibration curve was used to calculate the concentration of clofazimine in the brain homogenate samples.

### Retinal Imaging Procedures

Animal preparation and imaging procedures have been previously described (Bell et al., 2015). Briefly, mice were anesthetized using an IP injection of sodium pentobarbital (68 mg/kg). Mydriasis was induced using a 0.5 μl of 0.5% tropicamide phenylephrine mixture. Topical anesthesia was induced using 0.5% proparacaine. Cornea hydration and ocular media opacities were minimized using frequent applications of hydrating drops and topical eye shields (Bell et al., 2014). Following the procedure, eyes were covered with puralube ointment. Mice recovered in a warmed Plexiglas chamber with supplemental oxygen.

Confocal scanning laser ophthalmoscope (cSLO) imaging was performed using an HRA2 system (Heidelberg Engineering, Inc). A wide-field objective (55°) was used to image the retina with the optic nerve disk centrally located within the image frame. Imaging modes of infrared reflectance (IR-cSLO) at 800 nm and blue peak autofluorescence (BAF-cSLO) at 488 nm were used to image the retina and vitreoretinal interface.

Spectral-domain optical coherence tomography was performed following cSLO to examine and compare the in-depth retinal morphology between treatment groups. Orthogonal B-scans (1000 a-scans/b-scan x 15 frames) were collected through the optic disk from the horizontal and vertical meridians. The 15 frames from each meridian were co-registered and averaged using ImageJ and StackReg and TurboReg Plugins (Schneider et al., 2012; Thevenaz et al., 1998).

## Author contributions

EEM-H, LAT-H, and JDL provided conceptualization and design; EEM-H, LAT-H, DJS, JTE, ES, MH, BP, SAS, RZ, JSH, TA, AB, and BB performed the experiments; EEM-H, LAT-H, DJS, JTE, JZ, RZ, BB, PL, BKJ, and JDL analyzed the data; EEM-H and JDL wrote the manuscript; JDL provided financial support; and all authors provided final approval of the manuscript.

## Acknowledgements

We thank the members of the Lathia laboratory and the Reizes laboratory for insightful discussion and constructive comments on the manuscript. We thank Kevin Stoltz for assistance setting up the screening platform, Joseph Gerow and Eric Schultz for flow cytometry assistance, Amanda Mendelsohn for the illustrations included in this manuscript, and Earl Poptic and Melanie Hoffner in the LRI Molecular Screening Core for assistance with small molecule screening. This work was funded by the National Institutes of Health (grant NS089641), the Cleveland Clinic VeloSano Bike Race, and Cleveland Clinic Innovations. The Lathia laboratory also receives funding from the National Institutes of Health (grants NS083629 and CA157948), a Distinguished Scientist Award from the Sontag Foundation, and the Case Comprehensive Cancer Center.

