## Supplementary Materials for "Development of a Cx46 targeting strategy for cancer stem cells"

### Supplemental Information

**Supplemental Figure 1, related to Figure 1. Glioblastoma CSCs express Cx46 mutants.** (**A-B**) CSCs from the patient-derived xenograft specimens T4121 and T387 were transfected with wildtype or mutant Cx46 and lysed for RNA 48 h later. qPCR was performed using Fast Sybr Green, and results were analyzed using the ΔΔCt method. Expression is normalized to GAPDH and is shown relative to the vector-transfected cells. n = 3 experiments performed in triplicate. All data points are shown, with the mean indicated by a horizontal line. (**C**) CSCs from the patient-derived xenograft specimen T387 were transfected with wildtype or mutant Cx46, and the number of cells was measured on days 0, 1, 3, 7, and 10 after plating using CellTiter-Glo. The values shown are relative to day 0. n = 3 experiments performed in triplicate. ** $p<0.01$ by two-way ANOVA compared to vector to test for significant differences between the curves. Data are represented as mean ± SEM. (**D**) Summary table showing the published effects of each mutant on GJ and hemichannel activity when co-expressed with wild-type Cx46 and our observed effects on CSC characteristics. **–** indicates no change was observed.

**Supplemental Figure 2, related to Figure 4. Clofazimine likely acts specifically to inhibit Cx46-mediated GJIC in CSCs.** (**A**) Example $IC_{50}$ curves for clofazimine (CFZ) in two GBM specimens. Cells were treated with increasing concentrations of clofazimine for 72 h, and cell number was determined using CellTiter-Glo. (**B**) T4121 CSCs were treated with 2 µM clofazimine for 6 hours and subjected to RNA sequencing in triplicate. Volcano plots showing the distribution of changes in transcripts by RNA sequencing. Genes with significant changes are shown as red dots. (**C**) Heatmap showing the RNA sequencing hits with the largest changes with clofazimine treatment compared to DMSO vehicle. Red indicates higher expression, while blue indicates lower expression within each gene. (**D**) T4121 CSCs and non-CSCs were lysed for RNA and subjected to qPCR for Kv1.3. Each sample was analyzed in triplicate. * $p<0.05$ by unpaired Student's t-test with Welch's correction compared to expression in CSCs. Data are represented as mean ± SEM.

**Supplemental Figure 3, related to Figure 5. Clofazimine does not cross the blood-brain barrier.** (**A**) Mass spectrometry chromatograms of brain homogenate of animals treated with 2.44 mg/kg clofazimine (CFZ) or vehicle. Animals were treated IP for 5 days, and brains were homogenized and subjected to mass spectrometry. Clofazimine dissolved in 80% methanol was used as an internal standard. n = 3 brains from each group (vehicle and CFZ), and the chromatogram for each brain is shown. The red arrow indicates the peak for clofazimine. (**B**) Brain sections of mice treated with clofazimine. Animals were treated IP with 25 mg/ml clofazimine or vehicle (not shown), and the drug was allowed to circulate for 10 minutes. The innate red fluorescence of clofazimine in the treated brain is compared to a brain incubated with the lowest concentration of clofazimine standard (16 µg/mL). Images are inverted to show fluorescence as black puncta. Bar, 1 mm. (**C**) Representative confocal scanning laser ophthalmoscope (cSLO) images from the retinas of three control and three clofazimine-treated mice. The central black disk is the optic nerve. Image FOV diameter = ~1.6 mm. Infrared (IR)-cSLO images show IR signal at a wavelength of 800 nm being reflected from immediately adjacent retinal pigment epithelial (RPE) cells and choroidal tissue structures. Blue-light fundus autofluorescence (BAF)-cSLO images show the autofluorescence (excitation 488 nm/emission 500-700 nm) spanning signal emerging from the RPE monolayer that is specific to age-related lipofuscin accumulation from daily photoreceptor outer segment phagocytosis. IR-cSLO images of the vitreoretinal interface show the superficial retinal vasculature and nerve fibers originating from the optic nerve. Any toxicity to the retina of clofazimine treatment would have elicited strong inflammatory responses at one or more of the locations that would have been easily observed by non-invasive cSLO imaging.

**Supplemental Table 1. Primer sequences used.**

| Primer Name | Sequence (5' – 3') |
|---|---|
| Cx46 L11S F | AGCTTTCTGGGAAGACTCTCAGAAAATGCACAGGAGCAC |
| Cx46 L11S R | GTGCTCCTGTGCATTTTCTGAGAGTCTTCCCAGAAAGCT |
| Cx46 T19M F | AATGCACAGGAGCACTCCATGGTCATCGGCAAGGTTTGG |
| Cx46 T19M R | CCAAACCTTGCCGATGACCATGGAGTGCTCCTGTGCATT |
| Cx46 C54A F | GAGCAGTCAGACTTCACCGCCAACACCCAGCAGCCGGGC |
| Cx46 C54A R | GCCCGGCTGCTGGGTGTTGGCGGTGAAGTCTGACTGCTC |
| Cx46 C61A F | AACACCCAGCAGCCGGGCGCCGAGAACGTCTGCTACGAC |
| Cx46 C61A R | GTCGTAGCAGACGTTCTCGGCGCCCGGCTGCTGGGTGTT |
| Cx46 C65A F | CCGGGCGCCGAGAACGTCGCCTACGACAGGGCCTTCCCC |
| Cx46 C65A R | GGGGAAGGCCCTGTCGTAGGCGACGTTCTCGGCGCCCGG |
| Cx46 C181A F | CTGAAGCCGCTCTACCGCGCCGACCGCTGGCCCTGCCCC |
| Cx46 C181A R | GGGGCAGGGCCAGCGGTCGGCGCGGTAGAGCGGCTTCAG |
| Cx46 C186A F | CGCGCCGACCGCTGGCCCGCCCCCAACACGGTGGACGCC |
| Cx46 C186A R | GGCGTCCACCGTGTTGGGGGCGGGCCAGCGGTCGGCGCG |
| Cx46 C192A F | GCCCCCAACACGGTGGACGCCTTCATCTCCAGGCCCACG |
| Cx46 C192A R | CGTGGGCCTGGAGATGAAGGCGTCCACCGTGTTGGGGGC |
| Kv1.3 F | CAAAACGGGCAATTCCACTG |
| Kv1.3 R | TGAGCACAGCATGTCACTTG |
| Cx46 F | TGCACAGGAGCACTCCA |
| Cx46 R | GCGTGGACACGAAGATGAT |

**Supplemental Figure S1**

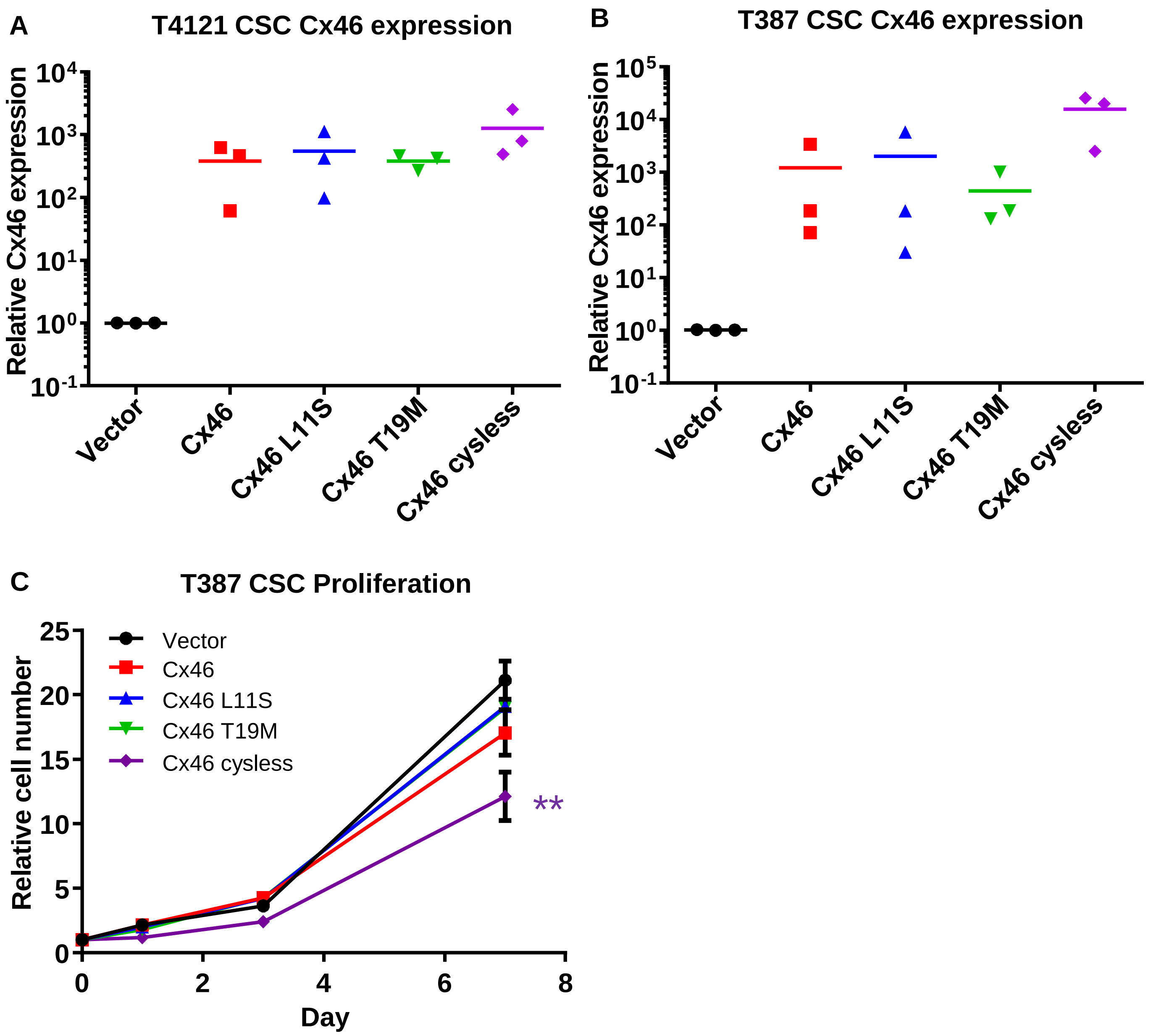


D

| Mutant | Reported effect on Cx46 function | | | Our observed phenotype in CSCs | | |
|---|---|---|---|---|---|---|
| | Hemi-channel | Channel | System | Growth | Survival | Self-renewal |
| L11S | ↓ | ↓ | Xenopus oocytes (Tong et al. Am J Physiol Cell Physiol 2013) | ↓ | -- | ↓ |
| T19M | ↑ | ↓ | Xenopus oocytes and HeLa cells (Tong et al. J Membrane Biol 2015) | -- | -- | -- |
| Cysless | -- | ↓↓↓ | Granulosa and MDCK cells (Tong et al. J Cell Sci.2007) | ↓↓↓ | ↓↓↓ | ↓↓↓ |

### Supplemental Figure S2

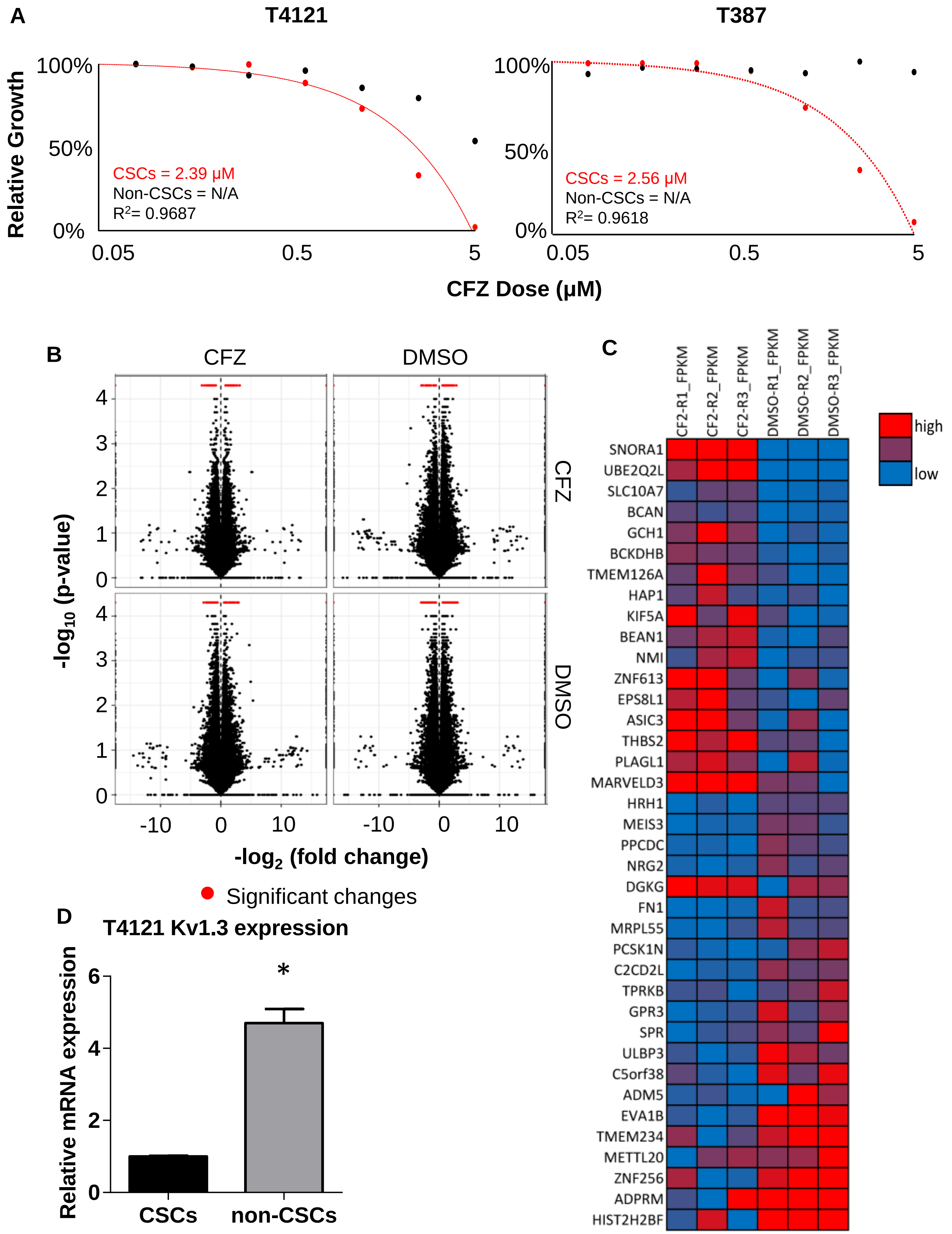

### Supplemental Figure S3

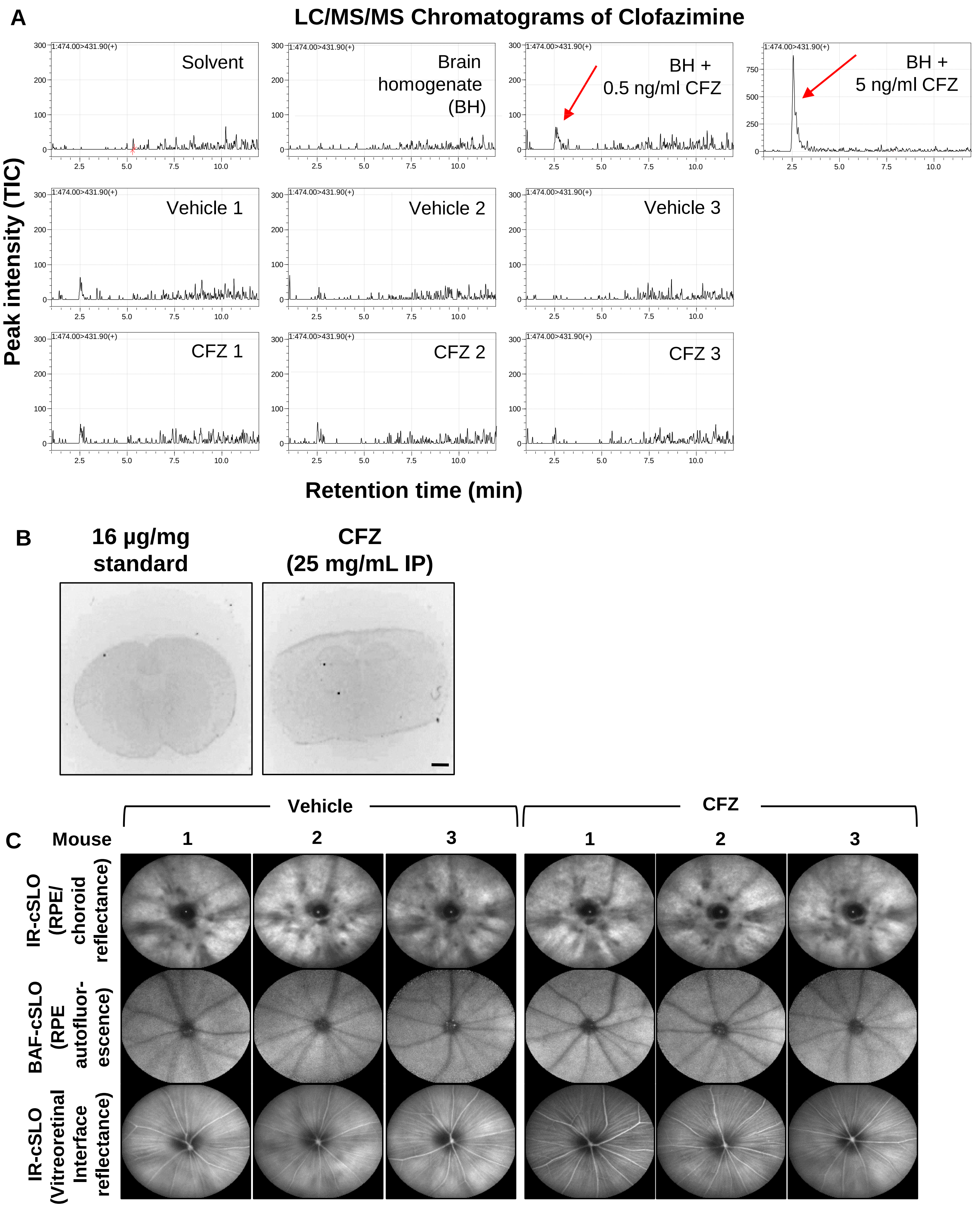
